# Different transcriptional responses to developmental versus short-term acclimation temperatures in *Pieris rapae*

**DOI:** 10.1101/2025.11.11.686846

**Authors:** Samantha L. Sturiale, Lorrie Le Yi He, Katherine H. Malinski, Christopher S. Willett, Joel G. Kingsolver, Leslie Ries, Peter A. Armbruster

## Abstract

To anticipate the role of thermal plasticity in evolutionary responses to climate change, it is critical to identify the molecular processes which underlie responses to temperature across different timescales. However, existing transcriptomic studies largely focus on responses to acute thermal stress or do not manipulate temperature across multiple timescales. We used RNA-sequencing to measure gene expression of *Pieris rapae* larvae exposed to a full factorial combination of non-stressful high and low temperatures across a long-term developmental timescale and a short-term acclimation timescale. This study design allowed us to separate genes associated with developmental thermal plasticity versus short-term acclimation responses, respectively. We observed that few genes were differentially expressed in response to both developmental temperature and short-term acclimation temperature, though there were some functional similarities across the two gene sets. This result suggests that the expression of different genes underlies thermal plasticity acting on different timescales, and thus these responses may evolve independently. Genes responsive to developmental temperature include those related to hormone activity and cold acclimation, while short-term acclimation temperature affected the expression of several cuticle protein genes. Both developmental and short-term acclimation temperature treatments affected the expression of genes involved in detoxification and protein folding. Finally, we identified a small subset of genes for which expression levels were dependent on the interaction between developmental and short-term acclimation temperature treatments, providing possible mechanisms by which developmental temperature may affect an organism’s capacity for acclimation responses later in life.

## INTRODUCTION

A key goal in ecology and evolutionary biology is to understand the role of phenotypic plasticity in the response of organisms to novel environmental conditions (Gibert et al., 2019; Levin, 1968; Levis and Pfennig, 2018; Nicoglou, 2015; Via et al., 1995; West-Eberhard, 2005). Plasticity in response to temperature (i.e., thermal plasticity) has received growing attention due to recent and ongoing climate change, as species are increasingly exposed to shifts in average temperatures, climatic variability, and seasonal timing (Coumou et al., 2013; Kodra et al., 2011; Kunkel et al., 2004; Linderholm, 2006; Menzel et al., 2006; Williams et al., 2015). As a result, numerous studies have investigated the fitness consequences of thermal plasticity to understand how it will influence the ability of ectothermic species to persist in novel climates (Gibbin et al., 2017; Kelly, 2019; Logan and Cox, 2020; Morley et al., 2019; Rodrigues and Beldade, 2020; Sandoval-Castillo et al., 2020; Sgrò et al., 2016).

Climate change alters temperature variability across multiple timescales, from heat waves and diurnal fluctuations to seasonal shifts and changes in mean annual temperatures (Intergovernmental Panel on Climate Change (IPCC), 2023). This variability presents a major challenge for understanding the fitness consequences of thermal plasticity because the phenotypic effects of temperature depend on (1) the magnitude, duration (timescale), and predictability of exposure; (2) the timing of the exposure in the individual’s life cycle; and (3) interactions across timescales (Kefford et al., 2022; Schulte et al., 2011; Stager et al., 2024). Temperatures experienced on relatively long timescales during early life stages can alter traits such as growth rate, resting metabolic rate, size, and thermal tolerance through developmental thermal plasticity (Beldade et al., 2011; Buckley and Kingsolver, 2021; Pottier et al., 2022). An alternative form of thermal plasticity called acclimation or sometimes ‘phenotypic flexibility’ (Piersma and Drent, 2003), can operate on either long-term (days to weeks) or short-term (minutes to hours) timescales throughout an organism’s lifetime, causing shifts in metabolism, behavior, physiology, and thermal tolerance (Armstrong et al., 2012; Clark et al., 2008; Goerge and Miles, 2024; Magozzi and Calosi, 2015; Teets et al., 2012). Importantly, the effects of acclimation are typically reversible, allowing an individual to repeatedly adjust their phenotype to novel environmental conditions, while developmental thermal plasticity results in stable changes to an individual’s phenotype (Schulte et al., 2011; Stager et al., 2024). Finally, there can be interactions across timescales, whereby the response to current temperature depends on the individual’s prior thermal environment (Alston et al., 2020; Beaman et al., 2016; Karl et al., 2012; Kingsolver et al., 2015; Stillwell and Fox, 2005).

While many studies quantify responses to temperature at just a single timescale, both developmental thermal plasticity and acclimation responses, as well as their interactions, are likely to be important to species’ responses to novel climatic conditions (Buckley and Kingsolver, 2021; Kefford et al., 2022). However, the roles of these distinct forms of thermal plasticity in species’ evolutionary responses to altered climate conditions will likely depend in part on the specific molecular processes underlying these different responses. For example, climate change is generally expected to increase environmental variability, which may favor acclimation over developmental plasticity if this variability occurs within the span of an organism’s lifetime, as acclimation allows an individual to repeatedly adjust their phenotype to their environment (Stager et al., 2024). However, if thermally plastic responses acting at different timescales are based on the same molecular processes or are mechanistically coupled due to interactions across timescales (Beaman et al., 2016), they might exhibit correlated responses to selection, impeding their independent evolutionary trajectories (Melo et al., 2016).

Despite the ubiquity of thermal plasticity across taxa as well as its potential importance to evolutionary responses to climate change, our understanding of the molecular processes which underlie developmental thermal plasticity versus acclimation is still limited. To investigate the molecular underpinnings of thermal plasticity, many researchers have used RNA-sequencing to detect genes whose expression changes in response to temperature and therefore potentially give rise to alternative phenotypes in target tissues (i.e., effector genes, (Lafuente and Beldade, 2019) (reviewed in (Oomen and Hutchings, 2017)). For example, exposure to heat shock is often associated with upregulation of heat shock proteins and detoxification pathways (Ashraf et al., 2022; Li et al., 2020; Shu et al., 2020; Zhang et al., 2023). However, existing studies have often focused on responses to stressful temperatures exposures of short duration (Ashraf et al., 2022; Li et al., 2020; Shu et al., 2020; Zhang et al., 2023) and/or do not manipulate temperature across multiple timescales (Chou et al., 2018; Chou et al., 2020; Des Marteaux et al., 2017; MacMillan et al., 2016). Additionally, of the few RNA-sequencing studies that do manipulate temperature across more than one timescale, most do not use the same temperature conditions for both exposure periods (Bellantuono et al., 2012; Lima and Willett, 2017; Metzger and Schulte, 2018; Scott and Johnston, 2012; Sørensen et al., 2016) (but see (Alston et al., 2020)). These limitations constrain both our ability to separate gene expression changes in response to non-stressful temperature exposures across timescales, and our understanding of the molecular mechanisms contributing to interactions across timescales. While there is some evidence that different genes underlie developmental thermal plasticity and short-term acclimation (Gerken et al., 2015), other studies have identified common transcriptional patterns or physiological processes associated with plastic responses to temperature at different timescales (Metzger and Schulte, 2018; Teets and Denlinger, 2013).

In this study, we sought to separate gene expression patterns associated with developmental thermal plasticity versus short-term acclimation responses in cabbage white (*Pieris rapae* (L.)) larvae. This small butterfly species is an important agricultural pest that has spread from its native range in Europe to multiple continents including North America (Harcourt et al., 1955; Richards, 1940; Ryan et al., 2019). Its thermal ecology has been well characterized (Jones and Ives, 1979; Kingsolver, 2000; Kingsolver et al., 2004; Richards, 1940), making it an excellent system with which to investigate the molecular mechanisms underlying distinct forms of thermal plasticity across timescales. We exposed *P. rapae* larvae to four combinations (two by two full factorial design) of non-stressful, diurnally fluctuating developmental and short-term acclimation temperature regimes then measured mortality, developmental time, and gene expression responses using RNA-sequencing (Fig. 1). Through this design, we were able to identify genes whose expression was affected by (1) developmental temperature, (2) short-term acclimation temperature, (3) both developmental and short-term acclimation temperature additively, and (4) the interaction of developmental and short-term acclimation temperature. This unique study design provides important insight into the molecular mechanisms responsible for developmental thermal plasticity and short-term acclimation responses as well as the interactions across these two timescales.

**Figure 1.**
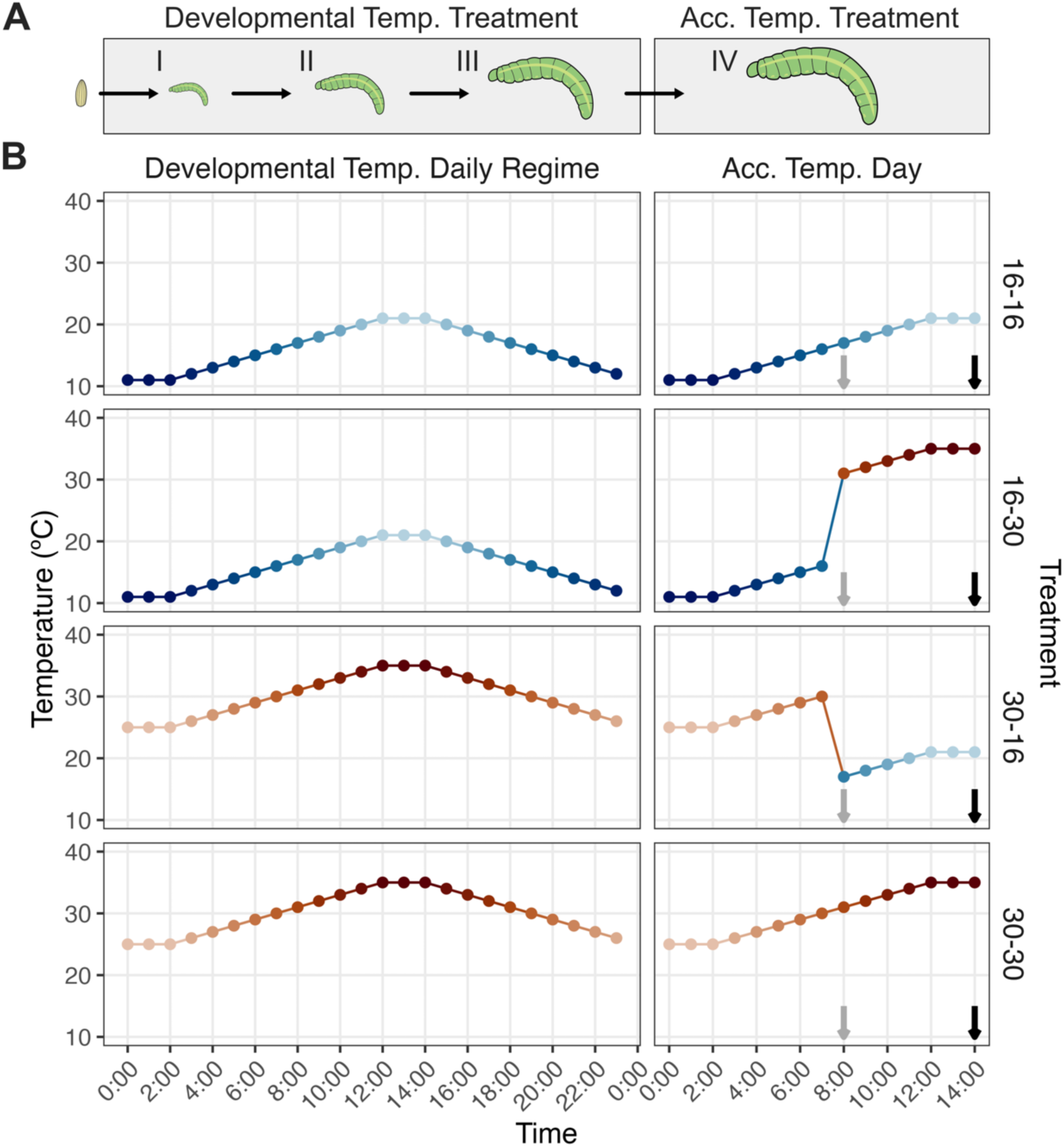
Schematic of the experimental design showing the life-history stages of developmental and short-term acclimation treatments (A) and the daily temperature profiles used for each treatment (B). (A) *Pieris rapae* larvae were exposed to their developmental temperature treatments from hatching through the third instar. On the day of molt to fourth instar, larvae were moved to their short-term acclimation temperature treatment (Acc. Temp. Treatment). (B) Larvae were exposed to a developmental temperature of either 16±5°C (top two panels) or 30±5°C (bottom two panels). For the short-term acclimation temperature treatment, larvae were either switched to the alternative temperature (middle two panels) or kept at the same temperature (top and bottom panels), resulting in four possible treatments (16-16, 16-30, 30-16, 30-30). The grey arrows indicate the time of day that the fourth instar larvae began their short-term acclimation temperature treatment, while the black arrows indicate the time of day that the short-term acclimation temperature treatments ended, and larvae were all snap-frozen for later RNA extractions. The colors of points and lines for all panels scale with temperature (warmest temperatures as dark red, coldest temperatures as dark blue).

## MATERIALS AND METHODS

### Field collections

Gravid females were collected in North Carolina in June 2022 (June 1^st^ – June 10^th^) from Perry-winkle Farm (Chapel Hill, NC) and kept under greenhouse conditions (∼24°C, 60-80% humidity, natural photocycle). Butterflies were kept in separate flight cages with 25% honey water as a food source and a collard plant (*Brassica oleracea*) as an oviposition stimulus. Plants were checked daily for new eggs, which were gently brushed onto a block of artificial wheat germ-based diet supplemented with collard powder as a feeding stimulus (Seiter and Kingsolver, 2013) in a large petri dish. Eggs were kept at room temperature (∼24-25°C) until hatching. Upon hatching, experimental larvae were transferred to diet blocks in individual petri dishes (100 x 15 mm, VWR #25384-342) and assigned a unique ID number. Five females generated eggs. Larvae from each of the five sibling groups were split approximately equally across the four treatments (described below) (range of 13-47 larvae per sibling group; Table S1).

### Developmental and short-term acclimation temperature regimes

This experiment was a two-by-two full factorial design with two temperature treatments and two timescales of temperature exposure (Fig. 1). The two temperature treatments consisted of exposure to 16°C or 30°C with ±5°C diurnal fluctuations. The first timescale of temperature exposure was from hatching until the day of molt to the fourth larval instar, here termed ‘developmental temperature’. The second timescale of temperature exposure was from 8:00am on the morning of molt to fourth instar until 2:00pm (six hours), here termed ‘short-term acclimation temperature’. These thermal regimes were chosen to maximize differences in physiology and life history without imposing thermal stress on the larvae based on previous work on this species (Jones and Ives, 1979; Kingsolver, 2000). Additionally, the 16±5°C and 30±5°C treatments are ecologically realistic, as they are consistent with average early spring versus mid-summer temperatures, respectively, in Chapel Hill, NC, where the population was collected (Supplementary File 1). Percival 36-VL environmental chambers were used to maintain a standard 14:10 L:D photoperiod and temperature conditions. Maximum and the minimum temperatures for each treatment were held for two hours daily, with linear ramping in between these two extremes. During their developmental temperature treatments, larvae were checked daily between 7:00 am – 8:00 am (Zeitgeiber time, ZT, 1-2) for molting or death to measure mortality rates and developmental time from hatch to fourth instar for each of the two developmental temperature treatments. Diet blocks were replaced every 48 hours.

On the day of molt (defined as when the headcap had fully slipped) to fourth instar, individuals were transferred to a short-term acclimation temperature of 30±5°C or 16±5°C at 8:00 am (ZT 2), then removed at 2:00 pm (ZT 8) and promptly flash frozen in liquid nitrogen and stored at –80°C until RNA extractions (Fig. 1). The starting time of the short-term acclimation temperature period was chosen to coincide with the midpoint in the diurnal temperature ramping program to minimize the risk of shock in treatment groups that switched between different temperature regimes. Thus, we collected RNA from four total treatment groups with different developmental and short-term acclimation temperature combinations: 30-30, 30-16, 16-30, and 16-16 (Fig. 1).

### JcDNV viral testing

At the time of experimental rearing, several individuals exhibited a larval-pupal intermediate phenotype, possibly indicating a viral infection was present in the population. Preliminary tests identified that these malformed individuals were infected by *Junonia coenia densovirus* (JcDNV), a virus that is commonly found in wild populations of *P. rapae* and other butterfly species, but that is not necessarily lethal to its host (McKeegan, 2024). Nevertheless, to minimize potential variation in the transcriptomic data associated with infection, we used PCR to screen frozen individuals for JcDNV infection prior to pooling samples for RNA extractions, retaining only JcDNV negative individuals for RNA-sequencing. We did not test for JcDNV infection in individuals which died during development, and thus we did not include infection status as a predictor variable in comparisons of mortality across developmental treatments. Infection status was however included as a predictor variable in comparisons of development time, and as a result we excluded a small subset of individuals who survived to fourth instar but were not tested for the virus (N = 15) from analyses of development time. A full description of the viral screening protocol is available in the Supplementary Materials.

### Statistical analysis of mortality rates and larval development time

All statistical analyses were conducted in R (version 4.3.2) (R Core Team, 2020). To compare mortality rates across the two developmental temperature treatments, we used a generalized linear mixed effects model with a binomial family distribution (glmer function in lme4 package version 1.1-35.5; (Bates et al., 2007)). In this model, developmental temperature was included as a fixed effect and maternal ID as a random effect. To compare days from hatch to fourth instar across the two developmental temperature treatments, we used a linear mixed effects model (lmer function in lme4 package) with developmental temperature and infection status as fixed effects and maternal ID as a random effect. Development time was square root transformed to meet the assumption of normality of residuals. The interaction between developmental temperature and infection status was included in the initial model but removed when found to be non-significant (*p* > 0.2) using a chi-squared test (Anova function in car package version 3.1-3 (Fox and Weisberg, 2019)). The life history and infection status data is available in supplementary file 1. The relevant coding scripts for analysis and visualization of the data can be found on the following GitHub repository: https://github.com/samstur/P_rapae_RNASeq/blob/main/MasterNotes.md.

### Tissue dissections and RNA extractions

We dissected out the muscle, fat body, and epidermal tissue (pooled tissues) from each caterpillar to use for RNA extractions because we were interested in understanding the effects of developmental and short-term acclimation temperature treatments on processes such as growth as well as energy storage and utilization (Arrese and Soulages, 2010). Specifically, for each caterpillar we cut the cuticle down the ventral side, removed the head along with the attached midgut, then collected the muscle, fat body, and epidermal tissue in the abdomen while disposing of the outer cuticle. Each biological replicate contained abdominal tissue from three to four caterpillars of the same developmental and short-term acclimation treatments. To minimize the effect of family, we combined tissues from caterpillars of two to four families for each biological replicate (Table S2). Combined tissues for each biological replicate were placed in 1 ml of RNAlater solution (Invitrogen, ThermoFisher Scientific, AM7020, Walthamm, MA, USA) and kept at 4°C overnight. The next day, each sample was transferred to a glass mortar, homogenized in 1 ml TRI Reagent (93289, Sigma-Aldrich, St Louis, MO, USA), then utilized for RNA extractions according to the manufacturer’s instructions. All RNA samples were then DNAase-treated using a TURBO DNA-free Kit (Invitrogen, ThermoFisher Scientific, AM1907, Walthamm, MA, USA). Four biological replicates for each of the four treatments (two developmental temperatures x two short-term acclimation temperatures) were chosen based on quality and quantity of RNA, which was measured on an RNA chip (Bioanalyzer 2100, Agilent Technologies, Santa Clara, CA, USA). These 16 total RNA samples were then used for cDNA library preparation and RNA sequencing at the Maryland Institute of Genome Science facility. All libraries were sequenced on a single lane of an Illumina NovaSeq 6000 instrument (150bp paired-end reads, strand-specific libraries). The RNA reads are available in NCBI’s short read archive (SRA) under BioProject accession PRJNA1258715.

### Bioinformatics processing and analyses

The details of the bioinformatics workflow and relevant coding scripts can be found on the following GitHub repository: https://github.com/samstur/P_rapae_RNASeq/blob/main/MasterNotes.md. Briefly, reads were trimmed for quality and to remove adapters using Trimmomatic (version 0.39) (Bolger et al., 2014) followed by quality control visualization by FastQC (version 0.11.9). We performed a two-pass mapping of all surviving reads to the latest genome assembly available on NCBI for *Pieris rapae* (GCF_905147795.1 aka ilPieRapa1.1) using STAR (version 2.7.1a) (Dobin et al., 2013). Then, HTSeq (version 0.13.5) (Anders et al., 2015) was used to count mapped reads for each gene. We removed genes with less than 10 reads total summed across all 16 samples before conducting differential expression analysis as described below.

Data visualization and differential expression analysis was performed in R (version 4.4.1). To visualize mRNA expression differences across sample types, we first performed principal components analysis (plotPCA function of DESeq2 package, version 1.44.0, (Love et al., 2014)) using variance stabilizing transformed read counts (varianceStabilizingTransformation function in DESeq2 package) for the 500 genes with the most variable expression across samples (the default for the plotPCA function). For differential expression analysis, the DESeq2 package was used to normalize counts for size factors and gene-wise dispersion estimates. The resulting normalized counts were then modeled using a negative binomial distribution and we performed a Wald test to compare the expression of each gene across treatments. The ashr package (version 2.2-63) (Stephens, 2017) was used to perform fold change shrinkage. We implemented two different model designs in our differential expression analysis with DESeq2. First, to obtain a list of genes which showed a significant effect of the interaction between developmental temperature and short-term acclimation temperature, we ran our differential expression analysis with a two-factor design with an interaction (developmental temperature + short-term acclimation temperature + developmental temperature:short-term acclimation temperature). We considered genes with a Benjamini–Hochberg false discovery rate adjusted *p*-value < 0.05 for the interaction term to be significantly affected by the interaction. These genes were not included in the second model and were analyzed separately. Second, we reran our analysis without the interaction (a design of developmental temperature + short-term acclimation temperature) to obtain a list of genes which exhibited a significant effect of developmental temperature and a list of genes which exhibited a significant effect of short-term acclimation temperature (adjusted p-value < 0.05, and |log_2_ fold change| > 1). For both developmental and short-term acclimation temperature factors, the lower temperature (16°C) group was assigned as the reference. As a result, a positive log_2_ fold change indicates higher expression at the higher temperature while a negative log_2_ fold change indicates higher expression at lower temperatures for both developmental temperature and short-term acclimation temperature treatments.

To examine groups of genes that were overrepresented in the differentially expressed (DE) gene lists for developmental temperature, short-term acclimation temperature, and their interaction, we performed overrepresentation tests using both gene ontology (GO) terms and KEGG pathways (enricher and enrichKEGG functions in the clusterProfiler package, version 4.12.6 (Yu et al., 2012)). To obtain GO annotations for *P. rapae* genes, we used the GO annotation file (.gaf) available for the *P. rapae* genome (GCF_905147795.1 aka ilPieRapa1.1) on NCBI, which was produced using InterProScan. Additionally, to improve our number of GO annotations, we combined the InterProScan annotations with an orthology-based approach. To do this, we assigned each *P. rapae* gene a best match to the *Drosophila melanogaster* proteome via blastx (only blastx matches with a bit-score > 45 and an e-value < 0.001 were used for GO term assignment). Each *P. rapae* gene then inherited the GO terms assigned to their best blastx *D. melanogaster* match. The background gene set list used for each overrepresentation test was the full gene output list associated with that term’s DESeq output (e.g., for the developmental temperature DEG list overrepresentation test, the background gene set was the full DESeq output for the developmental temperature term). GO terms and KEGG pathways were considered overrepresented if they had a Benjamini–Hochberg false discovery rate adjusted *p*-value < 0.05. To summarize overrepresentation results in our figures by removing redundant GO terms, we used REVIGO (Supek et al., 2011). The REVIGO program was run with the following settings: 0.5 similarity, the default semantic similarity measure (SimRel), the whole UniProt database, and included adjusted *p*-values from the overrepresentation test (with the setting “lower values is better”).

Lastly, we utilized hierarchical clustering to examine the different patterns of expression across samples in the list of genes that were DE in response to the interaction of developmental and short-term acclimation temperature. Specifically, we took normalized count data for each gene on this list (normalized for size factors and variance stabilizing transformed), scaled these counts by the mean and standard deviation across samples to obtain z-scores, and used these scaled counts to create a matrix of distances between each pair of genes. This matrix was then used to perform hierarchical clustering using the hclust function in the stats package (version 4.4.1) which grouped genes into clusters based on the degree of similarity in their patterns of expression across treatments. The number of clusters (eight) was chosen manually based a significant decrease in within-cluster dissimilarity, as visualized by an elbow plot.

## RESULTS

### The effects of developmental temperature on developmental time and mortality

There was no significant difference in larval mortality between developmental temperature treatments (Fig. 2A; χ²_1,167_ = 0.073, *p* = 0.787). Larval development time from hatch to fourth instar was significantly longer in the 16°C developmental treatment compared to the 30°C developmental treatment (Fig. 2B; χ²_1,107_ = 1581.314, *p* < 0.001). 24.5% of surviving larvae in the 16°C developmental treatment tested positive for JcDNV infection compared to 35.6% of larvae reared at 30°C. Larval development time was unaffected by JcDNV infection status (χ²_1,107_ = 0.006, *p* = 0.941) and the interaction between developmental temperature and JcDNV infection status was non-significant (*p* > 0.2) and so was removed from the final model.

**Figure 2.**
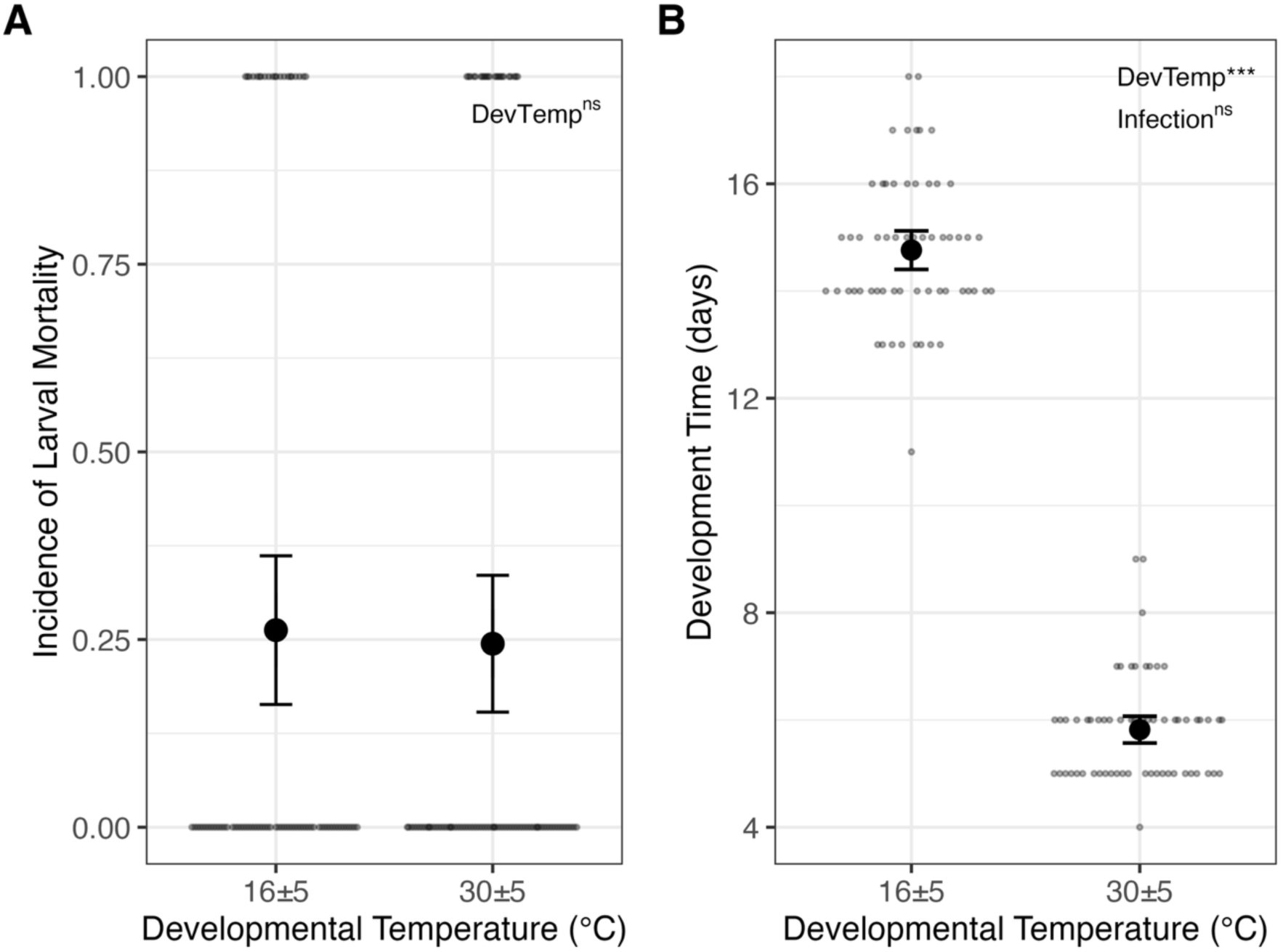
Developmental temperature did not affect larval mortality (A) but development at 30°C led to shorter development time from hatch to fourth instar compared to larvae reared at 16°C (B). Plots show means ± two standard errors for each treatment group as large points, while smaller lighter points indicate observations for individual larvae. JcDNV infection status did not affect development time. Sample sizes for (A) are N = 80 for 16°C group and N = 90 for 30°C group. Samples sizes for (B) are *N* = 59 for 16°C group and *N* = 61 for 30°C group. Effects of developmental temperature (DevTemp) and JcDNV infection (Infection) are indicated in the upper left of each panel, where ns indicates a non-significant effect and *** indicates a p < 0.001

### Quality control and principal components analysis of RNA-Sequencing data

Sequencing of mRNA produced between 44,434,324 – 66,032,118 reads per biological replicate. Across the 16 biological replicates, 96.2-96.6% of reads survived quality trimming and 83-89% of these trimmed reads aligned uniquely to the genome. Lastly, 85-91% of aligned reads were assigned to genes (Table S2).

The principal components analysis plot shows approximate clustering of the four treatment groups (Fig. S1). Specifically, samples clustered coarsely according to developmental temperature treatment along PC1, which explains 35% of variance, and separated more clearly by short-term acclimation temperature along PC2, which explains 19% of variance.

### The effect of developmental temperature on gene expression

In total, 221 genes were differentially expressed (DE) in response to developmental temperature treatment (Fig. 3, Supplementary File 2). Of these, 190 exhibited greater expression at the higher developmental temperature (positive log_2_ fold-change) while 31 exhibited greater expression at the lower developmental temperature (negative log_2_ fold-change).

**Figure 3.**
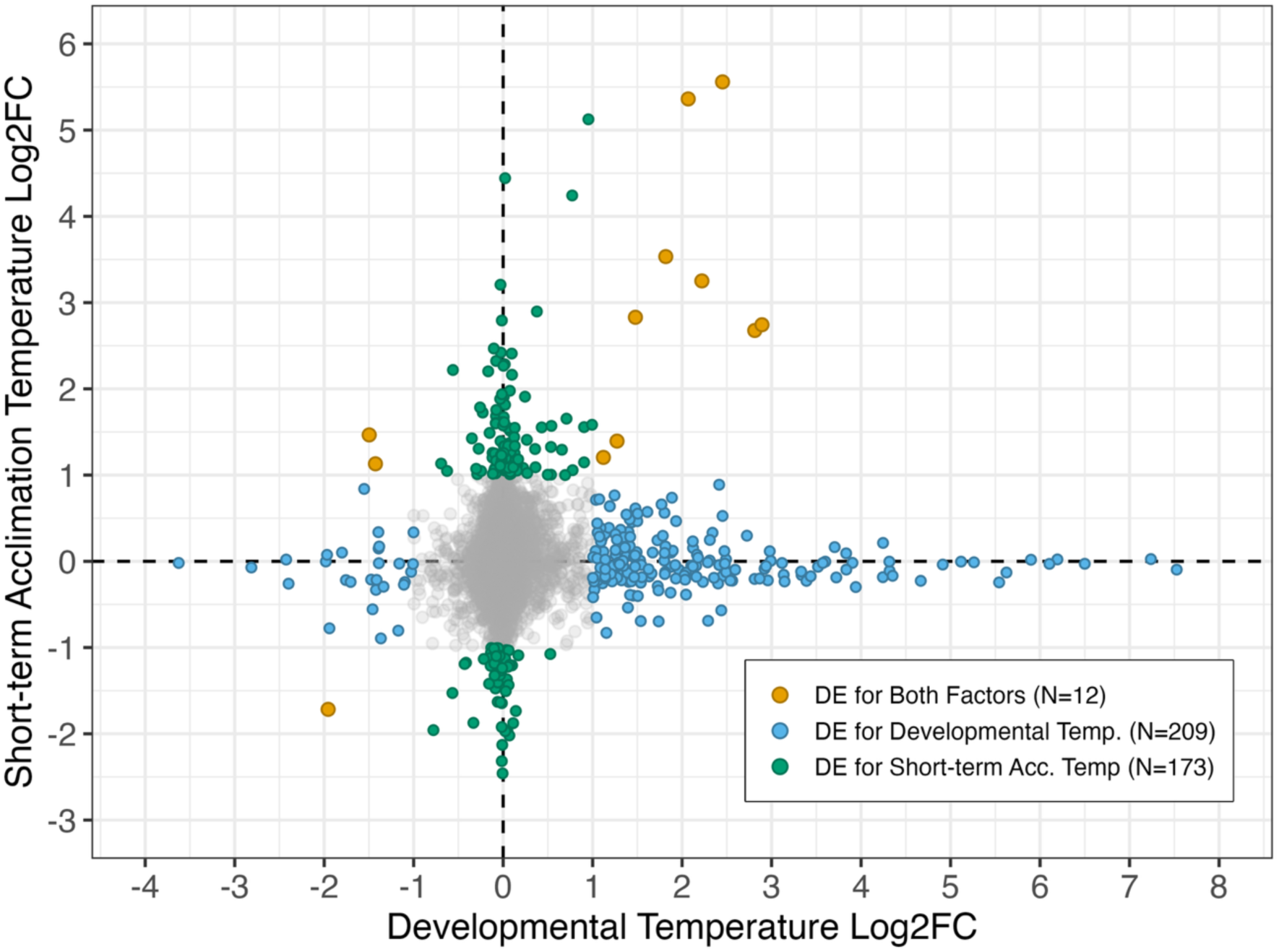
There is little overlap in genes differentially expressed in response to developmental temperature versus short-term acclimation temperature. Each point represents a gene, the color indicates whether the gene was significantly differentially expressed (DE) in response to only developmental temperature (blue), only short-term acclimation temperature (green), both factors (yellow), or neither factor (grey). The position of each point on the x– and y-axis indicates log_2_ fold-change. Genes which exhibited a significant interaction between developmental temperature and short-term acclimation temperature (*N* = 51) are not included in this plot. On both axes, a positive log_2_ fold change (log_2_FC) value indicates upregulation at higher temperature while a negative log_2_FC indicates upregulation at lower temperature.

**Figure 4.**
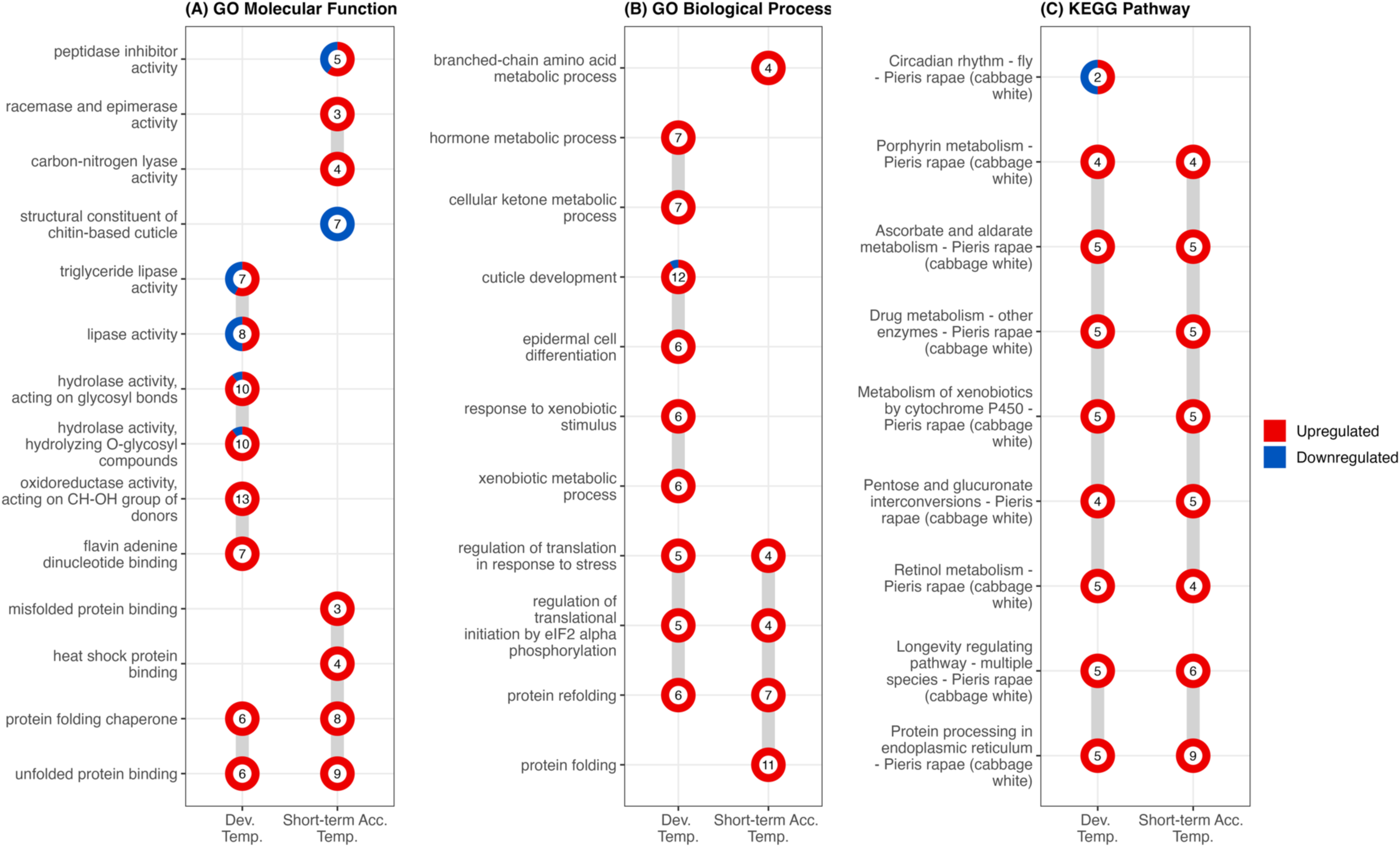
Enrichment of GO molecular functions (A), GO biological processes (B), and KEGG pathways (C) indicate some functional overlap in the genes differentially expressed in response to developmental temperature (Dev. Temp.) and short-term acclimation temperature (Short-term Acc. Temp.). For each enrichment term, the number of differentially expressed genes associated with that term is shown inside the circle, while the color of the outline indicates the relative proportion of associated differentially expressed genes that were upregulated (red) versus downregulated (blue) in response to higher developmental or short-term acclimation temperature. The grey lines connect related GO or KEGG terms (i.e., terms for which the list of associated differentially expressed genes overlap). For GO enrichment results, redundant parent/child terms are not shown, but a full list of enriched terms is available in Supplementary File 2.

Several GO molecular functions related to energy usage were uniquely enriched in the list of genes that were DE in response to developmental temperature including *triglyceride lipase activity* (*p*-adj = 0.003)*, hydrolase activity hydrolyzing O-glycosyl compounds* (*p*-adj = 0.002), and *oxidoreductase activity acting on CH-OH group of donors* (*p*-adj = 0.002) (Fig. 4A). We also observed that *protein folding chaperone* and *unfolded protein binding* molecular function terms were also significantly enriched, largely driven by heat shock proteins.

For GO biological processes, we observed overrepresentation in terms related to hormone regulation (*hormone metabolic process, p*-adj = 0.040), epidermal development (*epidermal cell differentiation, p*-adj = 0.007), and protein folding (*regulation of translation in response to stress*, *p*-adj < 0.001; *regulation of translational initiation by elF2 alpha phosphorylation, p*-adj < 0.001; and *protein refolding, p*-adj < 0.001) (Fig. 4B). The GO biological process *xenobiotic metabolic process* was also found to be overrepresented (*p*-adj = 0.003).

Lastly, the following KEGG pathways were overrepresented in the developmental temperature DE list: *circadian rhythm* (p-adj = 0.027), *porphyrin metabolism* (p-adj = 0.008)*, ascorbate and aldarate metabolism* (p-adj = 0.002)*, drug metabolism* (p-adj = 0.003)*, metabolism of xenobiotics by cytochrome p450* (p-adj = 0.007)*, pentose and glucuronate interconversions* (p-adj = 0.008)*, retinol metabolism* (p-adj = 0.002)*, longevity regulating pathway* (p-adj = 0.003)*, and protein processing in the endoplasmic reticulum* (p-adj = 0.031) (Fig. 4C).

Full enrichment results are available in the supplementary materials (Supplementary File 2).

### The effect of short-term acclimation temperature on gene expression

A total of 185 genes were DE in response to short-term acclimation temperature treatment. Of these, 131 exhibited greater expression at the higher temperature (positive log_2_ fold-change) while 54 exhibited greater expression at the lower temperature (negative log_2_ fold-change) (Fig. 3, Supplementary File 2).

Genes that were DE in response to short-term acclimation temperature were enriched for several molecular functions related to protein folding (Fig. 4A). Additionally, multiple terms related to amino acid metabolism were enriched (e.g., *peptidase inhibitor activity,* p-adj = 0.048; *racemase and epimerase activity,* p-adj = 0.048; and *carbon-nitrogen lyase activity,* p-adj = 0.007). Lastly, we observed enrichment of the molecular function term *structural constituent of chitin-based cuticle* (p-adj = 0.048), with all 7 of the DEGs in this group exhibiting downregulation (higher expression at the lower short-term acclimation temperature treatment).

We also found evidence for enrichment of GO biological functions related to protein folding and amino acid metabolism (Fig. 4B).

Finally, the following KEGG pathways were overrepresented in the short-term acclimation temperature DE list: *porphyrin metabolism* (p-adj = 0.040)*, ascorbate and aldarate metabolism* (p-adj = 0.010)*, drug metabolism* (p-adj = 0.040)*, metabolism of xenobiotics by cytochrome p450* (p-adj = 0.040)*, pentose and glucuronate interconversions* (p-adj = 0.013)*, retinol metabolism* (p-adj = 0.040)*, longevity regulating pathway* (p-adj = 0.048), and *protein processing in the endoplasmic reticulum* (p-adj = 0.003) (Fig. 4C). Notably, all of these KEGG pathways were also enriched in the developmental temperature DE list.

### Genes impacted by both developmental and short-term acclimation temperature treatments

Only 12 genes were significantly DE in response to both developmental and short-term acclimation temperature treatments, but not DE in response to the interaction (Fig. 3, Table 1). Of these genes, 10 exhibited the same direction of log_2_ fold-change for developmental and short-term acclimation temperature: nine genes exhibited greater expression at higher developmental and short-term acclimation temperatures; and one exhibited greater expression at lower developmental and short-term acclimation temperatures. The remaining two genes exhibited opposing patterns of log_2_ fold-change, with greater expression at lower developmental temperatures and at higher short-term acclimation temperatures (Fig. 3). Of the nine genes that were upregulated in response to both higher developmental and short-term acclimation temperature, five were putatively identified as heat shock proteins or other molecular chaperones (Table 1: b, d, e, g, h).

**Table 1.** 12 genes were significantly differentially expressed in response to both developmental temperature treatment and short-term acclimation temperature treatment. A positive log2FoldChange value indicates higher expression at a higher temperature while a negative log2FoldChange value indicates higher expression at a lower temperature.

| Gene | Gene Name | Developmental<br>Temperature<br>log<sub>2</sub>FoldChange | Developmental<br>Temperature<br>adjusted p-value | Short-term<br>Acclimation<br>Temperature<br>log<sub>2</sub>FoldChange | Short-term<br>Acclimation<br>Temperature<br>adjusted p-value |
| --- | --- | --- | --- | --- | --- |
| (a) LOC110992291 | uncharacterized<br>LOC110992291 | 1.273 | 1.26E-02 | 1.394 | 1.02E-03 |
| (b) LOC110993680 | protein lethal(2)essential for<br>life | 2.452 | 3.62E-06 | 5.559 | 1.13E-29 |
| (c) LOC110994457 | nuclear anchorage protein 1-like | 1.121 | 1.53E-02 | 1.205 | 1.61E-03 |
| (d) LOC110995990 | heat shock protein 68 | 1.478 | 7.53E-05 | 2.830 | 1.70E-17 |
| (e) LOC110996066 | alpha-crystallin A chain-like | 2.221 | 3.49E-07 | 3.251 | 1.30E-16 |
| (f) LOC110998413 | mucin-4 | -1.955 | 5.75E-08 | -1.717 | 3.91E-08 |
| (g) LOC110998985 | protein lethal(2)essential for<br>life-like | 2.068 | 1.33E-04 | 5.362 | 2.69E-26 |
| (h) LOC110999593 | protein lethal(2)essential for<br>life | 1.818 | 8.11E-04 | 3.533 | 4.51E-12 |
| (i) LOC111000288 | ankyrin repeat domain-<br>containing protein 50 | -1.427 | 9.79E-04 | 1.131 | 2.75E-04 |
| (j) LOC111003221 | transcription factor cwo | -1.497 | 4.86E-18 | 1.466 | 5.97E-20 |
| (k) LOC111004444 | pancreatic triacylglycerol<br>lipase | 2.812 | 2.52E-04 | 2.677 | 1.62E-05 |
| (l) LOC111004484 | maltase 2 | 2.892 | 1.16E-04 | 2.743 | 8.33E-06 |

### Genes impacted by the interaction of developmental and short-term acclimation temperature

A total of 51 genes were significantly differentially expressed in response to the interaction of developmental and short-term acclimation temperature (adjusted p-value < 0.05), with just 20 of these 51 genes exhibiting a strong interaction effect (|log2 fold-change| > 1) (Fig. 5, Supplementary File 2). For these 51 genes, the effect of short-term acclimation temperature on expression level depended on the developmental temperature the caterpillars had previously experienced. These genes exhibited a variety of expression patterns across samples (Fig. 5).

**Figure 5.**
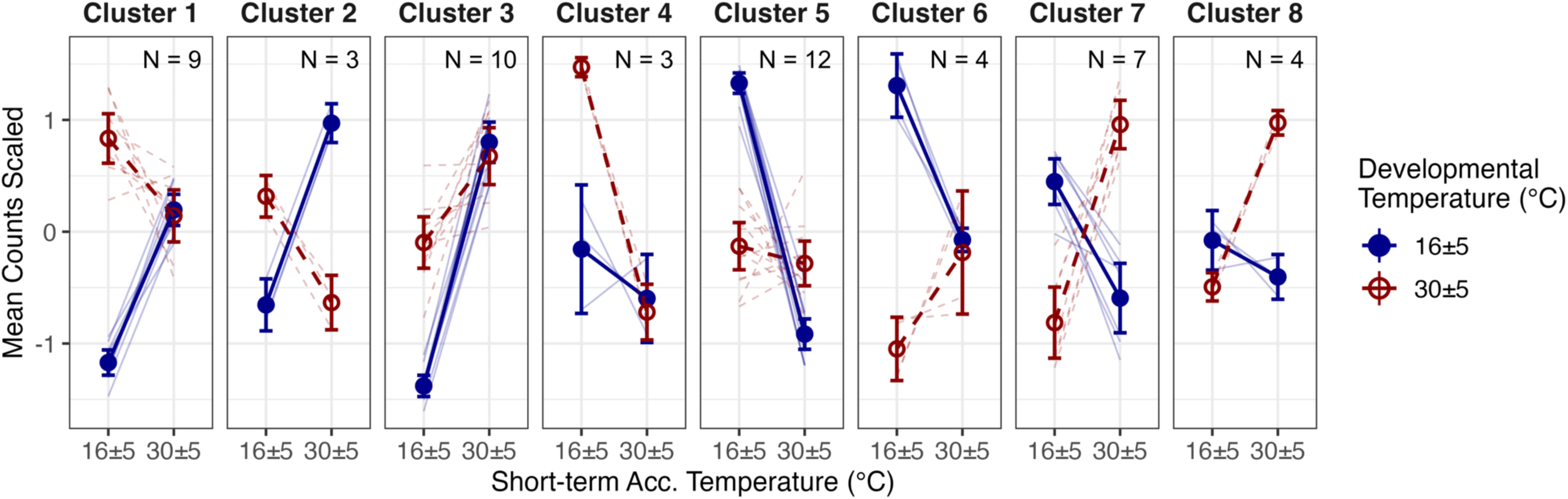
Expression patterns of the 51 genes which were significantly affected by the interaction of developmental temperature and short-term acclimation temperature. Genes are clustered based on their similarity in expression across treatment groups (see methods). Thin lines show the expression patterns (mean counts scaled across the four treatment groups) for each individual gene, while thicker lines and points show the mean expression of all genes within a given cluster (± two standard errors). Line-type and color indicate the developmental temperature group (dashed, red for 30°C and solid, blue for 16°C). The number of genes in each cluster is listed in the top right corner of each panel. For a list of genes in each cluster, see Supplementary File 2.

Genes that were significantly affected by the interaction of developmental and short-term acclimation temperature were enriched for GO molecular functions related to protein folding and RNA helicase activity. Similarly, enriched GO biological processes included those related to protein folding and muscle development, both of which were driven largely by *protein lethal(2)essential for life* genes. Finally, several KEGG pathways related to protein folding were enriched in the interaction DE gene set (Supplementary File 2).

## DISCUSSION

In this study, we performed a full-factorial manipulation of both developmental temperature (i.e., hatch to fourth instar larvae) and short-term acclimation temperature (six hours on the day of molt to fourth instar) to measure gene expression responses of *P. rapae* larvae. Unlike previous experiments which have often measured transcriptional responses to extreme temperature shocks, we utilized temperature regimes that were non-stressful – as evidenced by the high survival rates observed for both developmental temperature groups (Fig. 2A) – but resulted in large differences in life history (Fig. 2B). Additionally, this experimental design uniquely allowed us to identify genes whose response to short-term acclimation temperature depended on an individual’s prior developmental temperature to better understand how interactions between different types of thermal plasticity manifest at the transcriptomic level.

### Transcriptional responses to developmental versus short-term acclimation temperature were largely distinct

We observed that only 12 genes were differentially expressed (DE) in response to both developmental and short-term acclimation temperature (Fig. 3; Table 1). In contrast, 209 genes were DE in response to only developmental temperature, and 173 genes were DE in response to only short-term acclimation temperature (Fig. 3). At the functional level, the DE gene lists for developmental versus short-term acclimation temperature were largely enriched for different GO biological processes and GO molecular functions, though there was overlap in GO term enrichment related to protein folding/processing and in KEGG pathway enrichment related to detoxification (discussed further below) (Fig. 4, Supplementary File 2). This suggests that while individual genes associated with transcriptional responses to developmental versus short-term acclimation temperature were largely different, there is some similarity in the functions of these gene sets.

The finding that developmental temperature and short-term acclimation temperature triggered largely distinct transcriptional responses suggests that developmental thermal plasticity and short-term acclimation are facilitated by different sets of genes and thus could evolve independently. This interpretation is consistent with the results of a genome-wide association study (GWAS) in *Drosophila melanogaster* (Gerken et al., 2015). Gerken et al. (2015) identified distinct sets of candidate loci associated with the capacity for developmental plasticity versus rapid cold hardening, though similarly to our results, they observed functional similarities across the two sets of candidate genes. Further, while a study using RNA-sequencing in threespine stickleback, *Gasterosteus aculeatus,* found evidence for some shared mechanisms between developmental thermal plasticity and adult acclimation, the magnitude of overlap between these responses was relatively small (Metzger and Schulte, 2018).

Finally, several studies have observed cumulative effects of multiple acclimation treatments on thermal tolerance (e.g., long-term cold acclimation and rapid cold hardening), implying that they must operate through at least partially independent mechanisms (Colinet and Hoffmann, 2012; McDonald et al., 1997; Powell and Bale, 2005; Shintani and Ishikawa, 2007). Thus, our study contributes to growing evidence that there is modularity in the genes and mechanisms associated with plastic responses acting on different timescales.

Evidence for modularity or compartmentalization of plasticity-associated genes has also been observed in studies that compare plasticity of different traits in response to temperature. For example, Lafuente and colleagues found in their GWAS of *D. melanogaster* lines that there was little overlap in quantitative trait loci (QTLs) associated with variation in thermal plasticity for different body parts (i.e., pigmentation in thorax versus abdomen), for different pigmentation components (i.e., overall darkness versus pattern), and between temperature-driven pigmentation plasticity and size plasticity (Lafuente et al., 2018; Lafuente et al., 2024). The authors argue that their results conflict with the idea of ‘organism-wide plasticity QTLs’ that could jointly respond to evolutionary forces (Lafuente et al., 2024). Instead, modularity of plasticity genes across traits and across timescales may promote diversity in these responses by allowing them to respond to selection independently without the constraint of genetic covariation (McGlothlin and Ketterson, 2008; Melo et al., 2016). This modularity may also accelerate adaptation to novel environmental conditions that favor one form of plasticity over the other compared to a scenario where plastic responses are evolutionarily coupled. For example, greater environmental variability resulting from climate change may select for greater acclimation responses that confer increased flexibility relative to the irreversible effects of developmental plasticity (Stager et al., 2024).

In the following subsections, we discuss how several gene groups were impacted by developmental versus short-term acclimation temperature, largely focusing on genes found to be thermally responsive in past studies. Past transcriptomic experiments have often utilized short-term, extreme temperature treatments. Through our study, we were able to evaluate whether these gene groups also respond to more ecologically realistic, non-stressful temperature exposures of either short– or long-term duration.

### Genes related to hormone activity were affected predominately by developmental temperature

Hormones have long been hypothesized to facilitate developmental plasticity by serving as an intermediary between environmental signals and the expression of alternative phenotypes in target tissues (Beldade et al., 2011). Studies of insect polyphenisms largely support this hypothesis; polyphenisms are often coordinated in part by either ecdysteroids or juvenile hormone (JH), two important classes of insect hormones (Emlen and Nijhout, 2001; Oostra et al., 2014; Rountree and Nijhout, 1995; Simpson et al., 2011). Less is known, however, about the role of hormones in cases of developmental thermal plasticity that produce non-discrete (i.e., continuous) phenotypes. In our study, we show that a higher developmental temperature results in faster development (Fig. 2B), and similar to studies on polyphenisms, we find evidence that hormones may be involved in mediating this developmental thermal plasticity. This evidence includes the differential expression (DE) of 10 genes related to the activity of hormones such as ecdysteroids and JH in response to developmental temperature but not short-term acclimation temperature (Table S3), as well as the enrichment of the developmental temperature DE gene set for the GO term *hormone metabolic process*. In contrast, only three genes with hormone-related annotations were affected by short-term acclimation temperature, with no hormone related GO term enrichment, suggesting that this gene group is likely more important in mediating responses to temperature on longer timescales. Notably, two hormone receptor genes (LOC111004234 ‘*probable nuclear hormone receptor HR3*’ and LOC111004472 ‘*hormone receptor 4*’) were very strongly upregulated at higher developmental temperatures but not affected by short-term acclimation temperature. These receptors serve as critical regulators of insect development through their interactions with ecdysteroid hormones (Guo et al., 2015; Horner et al., 1995; King-Jones et al., 2005; Ou et al., 2011; Zhao et al., 2018), and our results implicate expression of these genes as a potential mechanism by which an individual’s life history is shaped by developmental thermal plasticity.

### Developmental and short-term acclimation temperature each affected different sets of genes involved in detoxification

As part of the transcriptional responses to both higher developmental and short-term acclimation temperature, we observed the upregulation of several classes of insect detoxification enzymes. However, different individual genes responded to developmental temperature versus short-term acclimation temperature (Table S3).

Specifically, five UDP-Glycosyltransferases (UGTs), four cytochrome p450s (CYPs), and one glutathione-S-transferase (GST) were DE in response to only developmental temperature, while three UGTs and three CYPs were DE in response to only short-term acclimation temperature. The differential expression of these genes also drove enrichment of several KEGG pathways related to detoxification (e.g., ‘*Drug metabolism – cytochrome P450’, ‘Metabolism of xenobiotics by cytochrome P450’*, etc.) in response to both developmental and short-term acclimation temperature (Supplementary File 2). CYPs and UGTs play an important role in the metabolism of both endogenous and xenobiotic compounds (Després et al., 2007; Kinareikina and Silivanova, 2024).

Furthermore, several transcriptomic studies have noted upregulation of CYPs and UGTs in response to acute thermal stress (Li et al., 2020; Liu et al., 2017; Shu et al., 2020; Tao et al., 2023). GSTs also often increase in expression following acute thermal stress (Ashraf et al., 2022; Liu et al., 2017; Shu et al., 2020), and these enzymes likely function to counteract the oxidative stress induced by stressful temperatures (Jia et al., 2011; Lalouette et al., 2011; Miao et al., 2020). Our finding that these detoxification enzymes change expression at relatively benign temperature conditions implies that they may be induced as part of an anticipatory response to modulate heat tolerance (as in (Des Marteaux et al., 2017; Toxopeus et al., 2019)) at both long and short-term timescales and/or that they mediate responses to temperature unrelated to thermal tolerance (e.g., steroid hormone biosynthesis).

### Genes related to cold acclimation and cold stress

To persist at cold temperatures, insects often increase production and/or transport of cryoprotectants to facilitate supercooling and alter the lipid composition of their plasma membranes to preserve fluidity (reviewed by (Teets et al., 2023)). Accordingly, we found that the most highly upregulated genes at lower developmental temperature included two trehalose/sugar transporters and two phospholipases (Table S3), with no effect of short-term acclimation temperature on either type of gene. These genes may act as part of a cold acclimation response in *P. rapae*, which is consistent with previous studies showing upregulation of trehalose transporters in insects (Toxopeus et al., 2019) and greater expression and/or activity of phospholipases in plants (Gustavsson and Sommarin, 2002; Wan et al., 2009) in response to cold acclimation conditions. Two other studies have observed upregulation of trehalose transporters after short-term cold stress in insects (Huang et al., 2017; Tao et al., 2023), indicating that these genes may act as both a preparatory response and a rapid defense under extreme conditions.

In response to short-term acclimation temperature, eight genes with annotations related to cuticle proteins were uniquely differentially expressed (seven out of eight were upregulated at low temperature), while only one cuticle protein gene was affected by developmental temperature (Table S3). We also observed enrichment for the GO molecular function ‘*structural constituent of chitin-based cuticle*’ for the short-term acclimation temperature DE gene set. Differential expression of cuticle proteins has been noted often in insect responses to short-term cold stress (Cui et al., 2017; Dennis et al., 2015; Dunning et al., 2014; Guo et al., 2018; Tao et al., 2023) and our study suggests that this gene group may also act on short-term timescales at relatively benign temperatures.

### Heat shock proteins and other genes related to protein folding

Heat shock proteins (HSPs) serve as molecular chaperones to prevent the aggregation of misfolded proteins (reviewed in (King and Macrae, 2015)) and are often upregulated following exposure to acute cold and heat shocks (Alston et al., 2020; Ashraf et al., 2022; Li et al., 2020; Sinclair et al., 2007), as well as during long-term or short-term thermal acclimation (Colinet and Hoffmann, 2012; Des Marteaux et al., 2019; Rosendale et al., 2022; Toxopeus et al., 2019). In our experiment, HSPs and other genes with annotations related to protein folding made up five out of 12 of the genes DE in response to both developmental temperature and short-term acclimation temperature, with all five showing greater expression at higher temperatures (Table 1: b, d, e, g, h). Additionally, we observed the enrichment of GO terms and KEGG pathways related to protein folding/processing in response to both higher developmental and short-term acclimation temperatures (Fig. 4, Supplementary File 2). These results suggest that molecular chaperones may be an important mechanism underlying both developmental thermal plasticity as well as short-term acclimation responses. However, both the number of DE molecular chaperone genes and the magnitude of their upregulation were greater in response to short-term acclimation temperature than developmental temperature (*N* = 6 for developmental temperature; *N* = 11 for short-term acclimation temperature; Table S3). Thus, the expression of HSPs and other molecular chaperones appears to more closely track short-term, recent thermal conditions, which may be beneficial to fitness given the potential costs to development and fecundity associated with chronically high HSP levels (reviewed in (Sørensen et al., 2003)).

### Genes which exhibited an interactive effect of developmental and short-term acclimation temperature

We tested for interactive effects of developmental temperature and short-term acclimation temperature on genome-wide transcription levels in *P. rapae* larvae to investigate the mechanisms by which an individual’s developmental temperature may affect its capacity for acclimation responses later in life. Several past experiments have observed significant interactions between developmental temperature and adult acclimation temperature on thermal tolerance (Healy et al., 2019; Kellermann et al., 2017; Schaefer and Ryan, 2006; Slotsbo et al., 2016b; Willot et al., 2021) or life history traits (Beaman et al., 2016; Stillwell and Fox, 2005). However, few studies have investigated the molecular underpinnings of this interactive effect, and those which have often compare the expression of a subset of genes rather than using untargeted approaches (Colinet and Hoffmann 2012; Karl et al. 2012; Healy et al. 2019). Two exceptions include RNA-sequencing studies by Alston and colleagues (2020), which found that transcriptional responses to heat shock depended on rearing temperature conditions in *Manduca sexta*, and by Scott and Johnston (2012), which observed an interactive effect of embryonic temperature and adult cold acclimation temperature on the expression levels of 11 genes in zebrafish (*Danio rerio*). Consistent with these two studies, we found that 51 genes exhibited a significant interaction effect of developmental temperature and short-term acclimation temperature (p-adjusted < 0.05), with just 20 of these genes exhibiting a strong interaction effect (|log2 Fold-change| > 1) (Supplementary File 2, Fig. 5). This number of genes is low relative to the groups of genes identified as DE in response to developmental and short-term acclimation temperature (221 and 185, respectively) as well as the number of total genes in the *P. rapae* genome (13,631). This low number may indicate that the organismal-level effects of interactions between developmental thermal plasticity and acclimation responses are mediated by a small number of genes. The interaction genes exhibited a variety of expression patterns across samples (Fig. 5). The expression of some genes responded to short-term acclimation temperature in the same direction but by different magnitudes across the two developmental temperature groups (Fig. 5; clusters 3, 4, and 5), while the expression of other genes exhibited opposing responses to short-term acclimation temperature depending on the developmental temperature treatment, leading to crossing reaction norms (Fig. 5; clusters 1, 2, 6, 7, 8).

The list of interaction genes was enriched for several GO biological processes, molecular functions, and KEGG pathways related to protein folding which resulted from the differential expression of four HSPs (Supplementary File 2). Intriguingly, for all four of these HSPs, the magnitude of expression increase in response to higher short-term acclimation temperature was greater in individuals reared at 16°C relative to individuals reared at 30°C (Fig. 5; cluster 3). This result implies that development at 30°C sets a higher minimum level of HSP expression than development at 16°C, thus reducing the responsiveness of HSP expression to short-term acclimation temperature in the 30°C developmental temperature treatment group. This pattern of lower HSP responsiveness after developing at higher temperature was also observed in a targeted transcriptomic study on *Lycaena tityrus*, which found that butterflies reared at 27°C exhibited higher baseline *HSP70* expression levels than individuals reared at 20°C, but that both groups had similar peak levels of expression following rapid heat hardening (Karl et al., 2012). A similar mechanism has also been proposed to underlie genetically based differences in thermal tolerance. Specifically, a study comparing the transcriptomic responses of thermally sensitive and thermally resilient corals to heat stress found that several heat shock proteins were less up-regulated after heat shock by more thermally resilient corals, a pattern they refer to as “constitutive frontloading” (Barshis et al., 2013). Our results suggest that a comparable frontloading mechanism – where the responsiveness of stress tolerance genes to short-term acclimation temperature is dependent on prior developmental temperature – could underlie previous observations of developmental temperature modulating adult acclimation capacity in thermal tolerance (Healy et al., 2019; Kellermann et al., 2017; Schaefer and Ryan, 2006; Slotsbo et al., 2016a; Willot et al., 2021).

### Conclusions

We exposed *P. rapae* larvae to a full factorial combination of non-stressful developmental and short-term acclimation temperature treatments, allowing us to separate genes associated with developmental thermal plasticity versus short-term acclimation responses, respectively. Notably, we found minimal overlap in the genes that were differentially expressed in response to differences in developmental temperature versus differences in short-term acclimation temperature. This result suggests that different genes underlie plastic responses acting on different timescales, and thus these responses may evolve independently. Several of the gene groups identified as differentially expressed have been associated with insect responses to temperature in past studies, including those related to hormone activity, detoxification, cold acclimation and/or cold shock, and heat shock proteins. Finally, we identified 51 genes for which expression levels were dependent on the interaction between developmental and short-term acclimation temperature treatments, providing possible links to mechanisms by which developmental temperature may affect an organism’s capacity for acclimation responses later in life. Rather than viewing thermal plasticity as a single monolithic trait, our results support a growing consensus that plastic responses acting at different timescales constitute largely distinct processes capable of exhibiting independent evolutionary trajectories during ectothermic species’ responses to novel climatic conditions (Kefford et al., 2022; Stager et al., 2024).

## LIST OF SYMBOLS AND ABBREVIATIONS

ZT: Zeitgeiber time
JcDNV: Junonia coenia densovirus
GWAS: genome-wide association study
QTL: quantitative trait locus
DE: differentially expressed
UGT: UDP-Glycosyltransferase
CYP: cytochrome p450
GST: glutathione-S-transferase
HSP: heat shock protein

## ACKNOWLEDGEMENTS

We thank Cathy Jones and Mike Perry of Perry-winkle Farms for permission to collect *P. rapae* at their farms. We also thank Madison A. Milotte for assistance with field collections and Dr. Mara Heilig for her advice regarding RNA extraction protocols. Work was funded by National Science Foundation awards IOS-2128241 to LR, PAA, CSW, and JGK and DEB-2128244 to JGK.

## AUTHOR CONTRIBUTIONS

SLS = conceptualization, methodology, software, formal analysis, investigation, writing – original draft preparation, visualization; LLYH = resources, methodology, investigation, writing-review, and editing; KHM = conceptualization, methodology, writing-review and editing; CSW = conceptualization, methodology, writing-review and editing; JGK = conceptualization, methodology, resources, writing-review and editing, supervision, funding acquisition; LR = conceptualization, methodology, writing-review and editing, supervision, funding acquisition; PAA = conceptualization, methodology, resources, writing-review and editing, supervision, funding acquisition.

## DATA ACCESIBILITY STATEMENT

The life history data and the results of differential expression and enrichment analyses are available as supplementary data files 1-2. The RNA reads are available on NCBI under the following BioProject accession number: PRJNA1258715. The details of the bioinformatics workflow and coding scripts can be found on the following GitHub repository: https://github.com/samstur/P_rapae_RNASeq/blob/main/MasterNotes.md

