## Supplementary Materials for "Different transcriptional responses to developmental versus short-term acclimation temperatures in *Pieris rapae*"

**Running title:** Thermal plasticity across timescales

**Key words:** phenotypic plasticity, thermal plasticity, gene expression, developmental plasticity, acclimation

**Author information:** Samantha L. Sturiale^1,‡^, Lorrie Le Yi He^2^, Katherine H. Malinski^2^, Christopher S. Willett^2^, Joel G. Kingsolver^2^, Leslie Ries^1^, Peter A. Armbruster^1^

^1^Georgetown University. Washington, D.C., USA

^2^University of North Carolina. Chapel Hill, North Carolina, USA

This PDF file includes:

Supplementary Methods (page 2)

Supplementary Tables (pages 3 – 8)

Supplementary Figures (page 9)

Supplementary References (page 10)

**Supplementary Methods**

*PCR screening for JcDNV infection:*

To test individual caterpillars for *Junonia coenia densovirus* (JcDNV) infection, snap-frozen whole caterpillars maintained at -80 °C were placed on a glass microscope slide on a glass petri dish on dry ice. A razor was used to remove the posterior abdominal segments. A new razor and microscope slide was used for each individual caterpillar and the glass dish was wiped down with ethanol and RNase-Zap (Invitrogen, AM9780, Waltham, MA, United States) to avoid cross-contamination and minimize exposure of the samples to RNases. The remainder of the caterpillar was returned to -80 °C for later tissue dissections and RNA extractions, while the dissected abdominal segments were placed in 500 microliters of 1XPBS and homogenized using a plastic pestle (*SP Bel-Art*, CAT No. 19923001, Warminster, PA, United States). 50 microliters of 20% N-Lauroylsarcosine sodium salt (ThermoScientific, CAT No. 434370250, Waltham, MA, United States) and 12.5 microliters of proteinase K were then added, and after brief mixing via inversion, the sample was incubated at 62 °C for an hour. Following the incubation, DNA was extracted using the DNAeasy blood and tissue kit (Qiagen, CAT No. 69504, Germantown, MD, United States). For each DNA sample, we ran two PCR reactions: one with a positive control primer set that amplifies a universal arthropod 28s ribosomal gene (Nice et al. 2009) and one with a viral primer set that amplifies the VP4 capsid protein gene of JcDNV (Wang et al. 2013). The 28s ribosomal control primer sequences were 5′ TACCGTGAGGGAAAGTTGAAA 3′ and 5′-AGACTCCTTGGTCCGTGTTT-3′ ′, forward and reverse primers respectively. The viral primer sequences were: 5′-CGAAAATCCACAGATACAACCAAG-3′ and 5′-GGGTTAGTATTAGCATCAGTCGAAAAT-3′, forward and reverse primers respectively. PCR amplification was performed using a GoTaq DNA polymerase (Promega, Madison, WI, United States) and FailSafe 2X PCR PreMix E (Lucigen, Berlin, Germany). We also ran a positive control of isolated viral DNA and a negative control with Ambion nuclease-free water (Invitrogen AM9930, Waltham, MA, United States) with each set of PCR amplification runs. Each PCR sample contained 1 microliter of sample DNA (or water for negative controls), 1 microliter of each primer (10 micromolar), 12.5 microliters of master mix, 9.13 microliters of nuclease-free water, and 0.37 microliters of polymerase. PCR thermocycler conditions were as follows: 95°C for 5 minutes, 45 cycles (95°C for 10 seconds, 60°C for 15 seconds, 72°C for 15 seconds), 72°C for 2 minutes. PCR products were visualized on a 1% agarose (Sigma-Aldrich A9539-250G, St. Louis, MO, United States) gel alongside a 1 Kb Plus DNA ladder (Invitrogen, CAT No. 10787018, Waltham, MA, United States). A caterpillar was identified as viral positive if their DNA sample produced a gel band for both the positive control PCR reaction and the viral PCR reaction. A caterpillar was identified as viral negative if their DNA sample produced a band for only the positive control PCR reaction.

**Supplementary Tables**

| *Table S1* Numbers of larvae from each of the five sibling groups assigned to each of the four treatments. Treatment numbers are written as ‘developmental temperature – short-term acclimation temperature’. | | |
| --- | --- | --- |
| **Sibling Group ID** | **Treatment** | **N** |
| 2 | 16-16 | 3 |
| 2 | 16-30 | 3 |
| 2 | 30-16 | 3 |
| 2 | 30-30 | 4 |
| 4 | 16-16 | 11 |
| 4 | 16-30 | 9 |
| 4 | 30-16 | 14 |
| 4 | 30-30 | 12 |
| 8 | 16-16 | 7 |
| 8 | 16-30 | 6 |
| 8 | 30-16 | 6 |
| 8 | 30-30 | 7 |
| 14 | 16-16 | 11 |
| 14 | 16-30 | 13 |
| 14 | 30-16 | 12 |
| 14 | 30-30 | 11 |
| 16 | 16-16 | 9 |
| 16 | 16-30 | 8 |
| 16 | 30-16 | 9 |
| 16 | 30-30 | 12 |

| *Table S2* A summary of the individual caterpillars pooled (based on their ID number), raw RNA-sequencing reads produced, and number and percentage of reads surviving processing steps (e.g., quality trimming, genome alignment, gene assignment) for each of the 16 biological replicates used for RNA-sequencing. RNA sample names correspond to mRNA read files available on NCBI. The first number in each caterpillar ID indicates its sibling group. | | | | | | | | | | |
| --- | --- | --- | --- | --- | --- | --- | --- | --- | --- | --- |
| **RNA Sample Name** | **Library ID** | **Treatment** | **Caterpillar ID numbers** | **Raw Read Count** | **Number of Surviving Reads After Trimming** | **Proportion of Surviving Reads After Trimming** | **Number of Reads That Uniquely Aligned to Genome** | **Proportion of Reads That Uniquely Aligned to Genome** | **Number of Reads Assigned to Genes** | **Proportion of Reads Assigned to Genes** |
| 16F-2 | IL100301767 | 16-30 | 4-20, 14-32, 16-52, 8-98 | 45271746 | 43681337 | 0.965 | 36320413 | 0.831 | 32941212 | 0.907 |
| 16F-3 | IL100301768 | 16-30 | 4-87, 14-152, 16-123, 8-113 | 44434324 | 42958337 | 0.967 | 36734732 | 0.855 | 33040017 | 0.899 |
| 16F-4 | IL100301769 | 16-30 | 4-60, 8-99, 14-155, 16-124 | 60123808 | 58150110 | 0.967 | 51239866 | 0.881 | 46692720 | 0.911 |
| 16F-5 | IL100301770 | 16-30 | 8-96, 8-97, 4-56, 14-158 | 63334777 | 61212166 | 0.966 | 54307537 | 0.887 | 49128830 | 0.905 |
| 16C-2 | IL100301771 | 16-16 | 2-176, 16-126, 8-91, 14-147 | 52570241 | 50868615 | 0.968 | 45596162 | 0.896 | 41122797 | 0.902 |
| 16C-3 | IL100301772 | 16-16 | 2-177, 4-66, 14-148, 8-111 | 61851447 | 59761252 | 0.966 | 52595795 | 0.88 | 46966013 | 0.893 |
| 16C-4 | IL100301773 | 16-16 | 2-69, 4-58, 8-110, 16-129 | 63354111 | 61295952 | 0.968 | 54727940 | 0.893 | 49230492 | 0.900 |
| 16C-5 | IL100301774 | 16-16 | 4-173, 8-93, 4-162 | 54078038 | 52203069 | 0.965 | 46117414 | 0.883 | 40499253 | 0.878 |
| 30F-1 | IL100301775 | 30-16 | 4-59, 14-36, 16-47, 14-143 | 58696957 | 56447412 | 0.962 | 48975499 | 0.868 | 43532294 | 0.889 |
| 30F-2 | IL100301776 | 30-16 | 4-67, 14-80, 16-51, 14-144 | 51495288 | 49826958 | 0.968 | 43957065 | 0.882 | 39100955 | 0.890 |
| 30F-3 | IL100301777 | 30-16 | 4-166, 8-116, 14-138, 16-74 | 55861001 | 53961765 | 0.966 | 47958730 | 0.889 | 42728614 | 0.891 |
| 30F-4 | IL100301778 | 30-16 | 4-179, 8-108, 14-139, 16-118 | 55983570 | 54027111 | 0.965 | 47516528 | 0.879 | 40677174 | 0.856 |
| 30C-2 | IL100301779 | 30-30 | 2-187, 4-57, 14-131, 16-73 | 58812970 | 56815772 | 0.966 | 50131864 | 0.882 | 44545480 | 0.889 |
| 30C-3 | IL100301780 | 30-30 | 4-7, 8-101, 14-134, 16-75 | 62244271 | 60207741 | 0.967 | 52418057 | 0.871 | 47250345 | 0.901 |
| 30C-4 | IL100301781 | 30-30 | 4-10, 8-114, 14-136, 16-76 | 66032118 | 63883869 | 0.967 | 55609919 | 0.87 | 49986378 | 0.899 |
| 30C-5 | IL100301782 | 30-30 | 4-164, 16-165, 4-65 | 60651846 | 58588797 | 0.966 | 51601834 | 0.881 | 46943291 | 0.910 |

| *Table S3* List of differentially expressed genes of interest grouped according to putative biological function as discussed in the text. Only genes that were differentially expressed in response to developmental and/or short-term acclimation temperature are shown in the table. NA values indicate that a gene was not differentially expressed in response to that treatment. Genes are arranged by their gene group (shown on the left side of table). | | | | | | |
| --- | --- | --- | --- | --- | --- | --- |
|  | **Gene** | **Gene Name** | **Developmental Temperature log_2_FoldChange** | **Developmental Temperature padj** | **Short-term Acc. Temperature log_2_FoldChange** | **Short-term Acc. Temperature padj** |
| Protein Folding Genes^a^ | LOC110993516 | heat shock protein 68-like | NA | NA | 5.125 | 5.81E-64 |
|  | LOC110994968 | dnaJ protein homolog 1 | NA | NA | 1.011 | 9.30E-48 |
|  | LOC110995067 | protein SGT1 homolog | NA | NA | 1.408 | 2.86E-87 |
|  | LOC110997862 | endoplasmin homolog | NA | NA | 1.057 | 2.86E-11 |
|  | LOC111002795 | activator of 90 kDa heat shock protein ATPase homolog 1 | NA | NA | 1.010 | 1.33E-30 |
|  | LOC111004285 | heat shock protein 68 | NA | NA | 1.909 | 1.27E-26 |
|  | LOC110993680 | protein lethal(2)essential for life | 2.452 | 3.62E-06 | 5.559 | 1.13E-29 |
|  | LOC110996066 | alpha-crystallin A chain-like | 2.221 | 3.49E-07 | 3.251 | 1.30E-16 |
|  | LOC110998985 | protein lethal(2)essential for life-like | 2.068 | 1.33E-04 | 5.362 | 2.69E-26 |
|  | LOC110998972 | protein lethal(2)essential for life-like | 1.886 | 2.53E-16 | NA | NA |
|  | LOC110999593 | protein lethal(2)essential for life | 1.818 | 8.11E-04 | 3.533 | 4.51E-12 |
|  | LOC110995990 | heat shock protein 68 | 1.478 | 7.53E-05 | 2.830 | 1.70E-17 |
| Hormone-related Genes^b^ | LOC110994502 | cytochrome P450 CYP12A2 | NA | NA | -1.957 | 4.05E-69 |
|  | LOC111003594 | adipokinetic hormone/corazonin-related peptide receptor variant I-like | NA | NA | -1.212 | 5.73E-04 |
|  | LOC111004327 | juvenile hormone esterase | NA | NA | 1.197 | 1.69E-03 |
|  | LOC111004234 | probable nuclear hormone receptor HR3 | 6.104 | 1.16E-09 | NA | NA |
|  | LOC110999701 | basic juvenile hormone-suppressible protein 1 | 4.245 | 3.92E-04 | NA | NA |
|  | LOC111004472 | hormone receptor 4 | 3.600 | 3.51E-06 | NA | NA |
|  | LOC110995266 | protein takeout | 3.335 | 1.12E-04 | NA | NA |
|  | LOC110992273 | juvenile hormone acid O-methyltransferase | 2.340 | 2.42e-06 | NA | NA |
|  | LOC110992426 | protein takeout | 1.811 | 4.63e-05 | NA | NA |
|  | LOC110998276 | juvenile hormone esterase | 1.256 | 1.00E-04 | NA | NA |
|  | LOC110999916 | uncharacterized LOC110999916 | 1.016 | 5.51E-04 | NA | NA |
|  | LOC111001029 | adipokinetic hormone/corazonin-related peptide receptor variant I | -1.704 | 2.81E-03 | NA | NA |
|  | LOC111003596 | neuroendocrine convertase 1 | -1.801 | 2.38E-03 | NA | NA |
| Detoxification Genes^c^ | LOC110994502 | cytochrome P450 CYP12A2 | NA | NA | -1.957 | 4.05E-69 |
|  | LOC110996979 | cytochrome P450 6k1 | NA | NA | 1.614 | 1.60E-06 |
|  | LOC110999362 | UDP-glycosyltransferase UGT5 | NA | NA | 1.598 | 5.53E-10 |
|  | LOC111001139 | cytochrome P450 6B5 | NA | NA | 1.304 | 1.32E-03 |
|  | LOC111003288 | UDP-glucosyltransferase 2 | NA | NA | 1.184 | 4.34E-04 |
|  | LOC111003292 | UDP-glycosyltransferase UGT5 | NA | NA | 1.576 | 2.00E-06 |
|  | LOC110998875 | cytochrome P450 6a8 | 3.835 | 1.00E-04 | NA | NA |
|  | LOC110999246 | UDP-glucosyltransferase 2 | 2.120 | 4.15E-03 | NA | NA |
|  | LOC110997327 | probable cytochrome P450 301a1, mitochondrial | 2.034 | 2.18E-04 | NA | NA |
|  | LOC110992284 | probable cytochrome P450 49a1 | 1.876 | 2.38E-03 | NA | NA |
|  | LOC110999237 | UDP-glycosyltransferase UGT5 | 1.814 | 1.48E-11 | NA | NA |
|  | LOC111001053 | glutathione S-transferase E14-like | 1.804 | 5.66E-05 | NA | NA |
|  | LOC111003289 | UDP-glucosyltransferase 2 | 1.504 | 2.16E-03 | NA | NA |
|  | LOC111002355 | probable cytochrome P450 304a1 | 1.416 | 6.99E-05 | NA | NA |
|  | LOC110999367 | UDP-glycosyltransferase UGT5-like | 1.290 | 7.74E-04 | NA | NA |
|  | LOC110992970 | UDP-glucosyltransferase 2 | -1.160 | 2.57E-03 | NA | NA |
| Cuticular Protein Genes^d^ | LOC110993166 | pupal cuticle protein C1B | NA | NA | 1.078 | 1.66E-04 |
|  | LOC110999339 | ice-structuring glycoprotein-like | NA | NA | -1.970 | 1.55E-08 |
|  | LOC110999341 | larval/pupal rigid cuticle protein 66-like | NA | NA | -1.506 | 1.22E-05 |
|  | LOC111001674 | endocuticle structural glycoprotein ABD-5-like | NA | NA | -1.206 | 4.23E-06 |
|  | LOC111001794 | larval/pupal cuticle protein H1C-like | NA | NA | -1.128 | 7.03E-04 |
|  | LOC111001799 | pupal cuticle protein G1A-like | NA | NA | -1.201 | 2.96E-05 |
|  | LOC111004346 | larval cuticle protein A2B-like | NA | NA | -1.921 | 1.60E-06 |
|  | LOC111004394 | pupal cuticle protein-like | NA | NA | -1.223 | 1.63E-05 |
|  | LOC110997778 | pupal cuticle protein Edg-84A | 3.833 | 2.38E-04 | NA | NA |
| Cold Acclimation Related Genes^e^ | LOC111002761 | phospholipase A1 VesT1.02-like | -1.385 | 3.62E-03 | NA | NA |
|  | LOC111002314 | facilitated trehalose transporter Tret1-like | -1.941 | 2.02E-06 | NA | NA |
|  | LOC110999177 | sugar transporter ERD6-like 12 | -1.981 | 1.27E-04 | NA | NA |
|  | LOC110993189 | phospholipase A2-like | -3.626 | 1.43E-04 | NA | NA |
| ^a^ Protein folding genes were identified with the following search term on NCBI gene: "pieris rapae"[Organism]) AND "protein folding"[Gene Ontology]  ^b^ Hormone-related genes were identified by querying *P. rapae* genome annotations for the terms “hormone” and “endocrin” and combining this list with the results from the following search term on NCBI gene: "pieris rapae"[Organism]) AND hormone  ^c^ Detoxification genes were identified by querying *P. rapae* genome annotations for the terms “UGT”, “UDP-glucosyltransferase”, “glutathione S-transferase”, “cytochrome P450”, “catalase”, and “superoxide dismutase”  ^d^ Cuticular protein genes were identified by querying *P. rapae* genome annotations for the term “cutic” and combining this with the list of genes with the GO molecular function annotation “structural constituent of chitin-based cuticle”  ^e^ Cold acclimation related genes were identified by querying *P. rapae* genome annotations for the terms “trehalose”, “sugar transporter”, and “phospholipase” | | | | | | |

**Supplementary Figures**

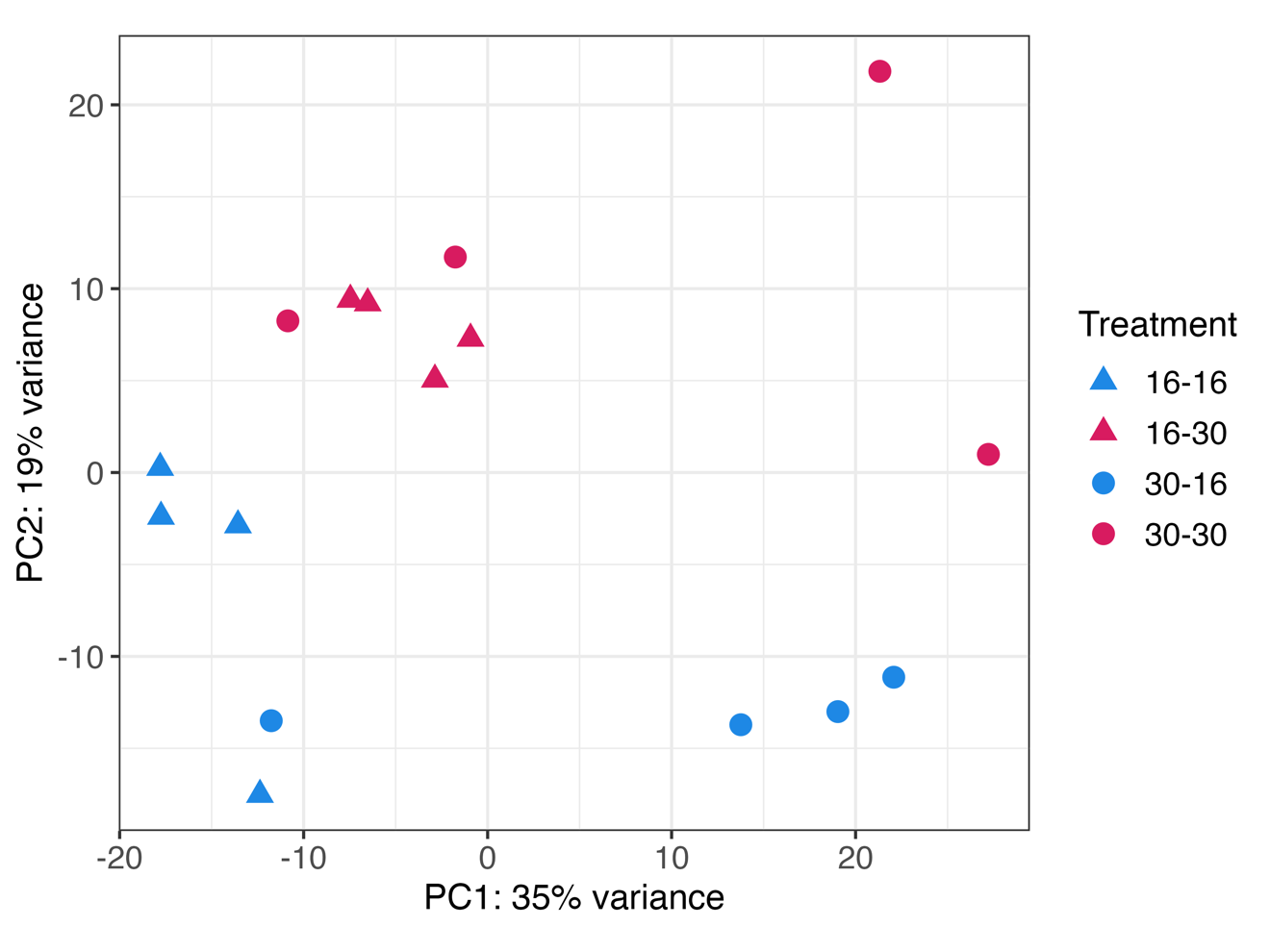

Supplementary Figure 1 Principal components analysis plot showing that the four temperature treatment groups roughly separate by developmental temperature on PC1 and by short-term acclimation temperature on PC2. The legend treatment labels are shown as “developmental temperature – short-term acclimation temperature”. The shape of the point indicates the developmental temperature treatment (triangle for 16°C and circle for 30°C), while the color indicates the short-term acclimation temperature treatment (blue for 16°C and pink for 30°C).
